# Ketone and glycolytic metabolism are key modulators of inflammation during neonatal sepsis

**DOI:** 10.1101/2025.08.15.670527

**Authors:** Björn Klabunde, Ole Bæk, Karoline Aasmul Olsen, Anna Hammerich Thysen, Margret Gudbrandsdottir, Katrin Laakmann, Kerstin Hoffmann, Bernd Schmeck, Anders Brunse, Nguyen Phuoc Long, Quoc Viet Le, Susanne Brix, Bo Chawes, Duc Ninh Nguyen

## Abstract

Neonatal sepsis is a life-threatening condition in preterm infants, primarily due to a dysregulated immunometabolic response to infection. Sepsis and infection mortality are associated with excessive glycolysis-induced inflammation, impaired mitochondrial oxidative phosphorylation (OXPHOS) and loss of disease tolerance. Reduced glucose intake can reverse these dysregulations, but it is unclear how the mechanistic control of glycolysis-OXPHOS balance drives defense strategies and infection outcomes. Here, in a preterm piglet model of neonatal sepsis, glycolysis inhibition with 2-deoxyglucose (2-DG) completely prevents acute infection mortality, reduces systemic inflammation and markers of liver injury, accompanied by enhanced mitochondrial metabolism and disease tolerance. Strikingly, this protection by 2-DG is conferred despite elevated blood glucose levels and higher bacterial burdens than the infected controls. Alternatively, partial replacement of glucose intake with the ketone beta-hydroxybutyrate (BHB) abolishes sepsis-related mortality via improving disease tolerance and clinical parameters. This intervention also shifts the hepatic transcriptome away from inflammatory signaling and towards mitochondrial metabolism. In macrophages *in vitro*, BHB also exerts anti-inflammatory effects independently of metabolic modulation via the HCAR2 receptor. Finally, data from a cohort of 700 infants confirm an association of plasma BHB levels and anti-inflammatory state. These findings demonstrate that metabolic reprogramming through glycolysis inhibition or ketone supplementation is a promising therapeutic strategy to enhance disease tolerance and improve sepsis outcomes in neonates.

## Introduction

Neonatal sepsis remains a significant cause of morbidity and mortality among preterm infants, particularly those born before 32 weeks of gestation ^1^. Late-onset sepsis, frequently caused by pathogens such as *Staphylococcus epidermidis*, affects up to 40% of preterm neonates and accounts for substantial neonatal mortality worldwide^1,2^. Their high sensitivity to infection is derived from the immature skin and gut barriers and frequent use of invasive medical interventions ^3^. However, the immune system of preterm neonates is also hyporesponsive during the early phase of infection, due to a unique metabolic phenotype, characterized by low energy reserves coupled with a high energy expenditure for growth and development^3^. Since defending against pathogens requires a large energy expenditure, initial immune responses are often muted. However, when the invading pathogen proliferates to a certain high threshold, a dysregulated and exaggerated immune response occurs, characterized by excessive immune activation causing profound organ damage^1,4^.

Effective defense against neonatal infections requires a delicate balance between two immune strategies: *disease resistance*, which aims to eliminate pathogens, and *disease tolerance*, which limits tissue damage without necessarily reducing pathogen load ^5^. Resistance is typically associated with a rapid inflammatory response and a metabolic shift toward aerobic glycolysis to support the energetic demands of activated immune cells^6,7^. In contrast, tolerance mechanisms often involve anti-inflammatory signaling and preservation of tissue integrity. Effective tolerance is supported by oxidative phosphorylation (OXPHOS) and efficient mitochondrial function ^8^. In neonatal sepsis, the balance of disease tolerance and resistance is frequently disrupted, and a loss of disease tolerance leads to organ dysfunction as the hallmark of sepsis. Metabolic disturbances occur during sepsis, including impaired mitochondrial function with reduced ATP production and elevated reactive oxygen species ^9,10^. In preterm infants, disturbances in glucose metabolism, such as hyperglycemia and hyperlactatemia, correlate with worse clinical outcomes and organ damage ^11–13^. This highlights that dysregulated energy metabolism and impaired defense strategies – particularly the loss of disease tolerance – are central to the pathophysiology of sepsis in neonates.

Preterm infants are commonly supplied with glucose-rich parenteral nutrition (PN) ^14^. This practice however increases the risks of hyperglycemia. Using a well-established piglet model of neonatal sepsis, we previously demonstrated that septic animals show elevated glycolytic activity in hepatic tissues, which is associated with systemic inflammation and loss of disease tolerance ^15^. We have further shown that high parenteral glucose supply accelerates hepatic glycolysis and systemic pro-inflammatory responses, increasing sepsis risk ^16–18^. In contrast, replacing glucose with galactose or glucogenic amino acids improved disease tolerance and infection survival ^18^. Despite these findings, the precise metabolic mechanisms underlying the protective effects of reduced hepatic glycolytic activity are unclear. It is also unknown whether the sepsis-preventive effect of glucose reduction predominantly stems from an altered host metabolism or from a growth-limiting effect on invading bacteria.

Likewise, boosting OXPHOS has been proposed as a metabolic intervention to reduce inflammation and improve disease outcomes^19,20^. Ketone bodies have recently emerged as metabolic supplements aimed at improving OXPHOS and mitochondrial activity^21,22^. Beta-hydroxybutyrate (BHB), the main ketone body produced during consumption of a ketogenic diet or prolonged fasting, was demonstrated to improve mitochondrial function in hyperexcited neurons^22^ and in a mouse model of transient ischemia^23^. In adults, higher blood plasma levels of BHB correlate with sepsis survival^24^. BHB also inhibits NLRP3 inflammasome activation during spinal cord injury and Alzheimer’s disease^24,25^

In the current study, we hypothesized that inhibition of glycolysis and activation of OXPHOS will improve disease tolerance and infection survival, and that BHB supplementation may be a novel, metabolism-centered treatment strategy during neonatal infections. We first investigated the hypothesis in animal experiments, examining the effect of 2-deoxy-glucose-mediated glycolytic inhibition and BHB treatment during experimental neonatal infection in piglets and then validated our findings in a human cohort of 700 children.

## Material and Methods

### Cultivation of *S. epidermidis*

*S. epidermidis* (isolated from a septic infant) was cultured in brain heart bouillon overnight at 37°C / 180 rpm. On the day of the infection, a day culture was inoculated from the overnight culture and incubated until the early logarithmic phase (OD600 = 1). Bacterial density was determined using a spectrophotometer. Bacteria were centrifuged and suspended in sterile saline to a final OD_600_ = 1, equivalent to 3 × 10^8^ CFU/ml. Piglets were inoculated with 10^9^ CFU/kg body weight by arterial infusion over 3 min. Control piglets were infused with a corresponding volume of sterile saline.

### Animal experiments

All animal studies and experimental procedures were approved by the Danish Animal Experiments Inspectorate under license number 2020-15-0201-00520. These approvals are in accordance with the EU Directive 2010/63, which governs the legislation for the use of animals in research. Crossbred preterm piglets (Landrace × Large White × Duroc) were delivered by cesarean section at day 106 of gestation (∼90% of term) from healthy sows. Immediately after birth, the piglets were transferred to a controlled intensive care unit, where they were housed individually in ventilated incubators preheated to 37°C with an initial oxygen supply of 1–2 L/min. If required, piglets were resuscitated using Doxapram and Flumazenil (0.1 mL/kg, intramuscular injection), tactile stimulation, and positive airway pressure ventilation until respiratory stability was achieved. Each piglet was equipped with a vascular catheter inserted into the dorsal aorta via the transected umbilical cord, allowing for parenteral nutrition (PN) administration, bacterial inoculation, and blood sampling. Approximately 2 h postpartum, all piglets were randomized into experimental groups based on birth weight and sex.

### Experimental groups and infections

Piglets received PN formulations differing in glucose and metabolic substrate composition and were challenged with live *S. epidermidis* (SE) or sterile saline (CON) via intraarterial infusion over 3 min. The following experimental conditions were used: Experiment 1 using PN with 10% glucose (14.4 g/kg/d), with or without 2-deoxy-D-glucose (2-DG; 750 mg/kg in 2.5 ml/kg PBS) (N=12 for infected groups; 2 for control groups). In experiment 2 piglets received PN with 10% glucose or 2.5% glucose (3.6 g/kg/d) + 2.5% β-hydroxybutyrate (BHB) (3.6 g/kg/d), or 2.5% glucose. Animals were monitored and cared for until euthanasia at 18 h post-infection or when humane endpoints were reached.

### Clinical monitoring and humane endpoints

Throughout the experiment, piglets were continuously observed for clinical signs of sepsis. Animals reaching predefined humane endpoints, including an arterial blood pH of ≤7.1, deep lethargy, discoloration, and tachypnea, were euthanized for tissue collection and categorized as non-survivors.

### Blood sampling and analysis

Blood samples were collected at 3, 6, 12 h post-inoculation and at euthanasia via the arterial catheter for blood gas analysis, hematology, and plasma storage for cytokine measurements. Additionally at euthanasia, serum and plasma samples were obtained for biochemical and metabolic analyses. To determine bacterial counts in blood, jugular vein punctures were performed at 3-and 6-h post-infection and at euthanasia. Blood samples collected at 3, 6, 12 h post-infection and at euthanasia underwent routine blood gas analysis using the GEM Premier 3000 (Instrumentation Laboratory, USA). Hematological assessments were performed using the ADVIA 2120i Hematology System (Siemens, Germany).

For cytokine analysis, plasma concentrations of TNF-α, IL-6, and IL-10 were quantified using porcine-specific DuoSet enzyme-linked immunosorbent assays (ELISA) (R&D Systems, USA) according to manufacturer’s instructions. Serum biochemistry evaluations, including liver and kidney function markers, were conducted using the ADVIA 1800 Chemistry System (Siemens, Germany).

### Euthanasia and tissue collection

At humane endpoints or scheduled euthanasia times, all piglets were deeply anesthetized with a Zoletil mixture (0.1 ml/kg), which included Zoletil 50 (125 mg tiletamine, 125 mg zolazepam), xylazine (6.25 ml xylazine 20 mg/ml), ketamine (1.25 ml ketamine 100 mg/ml), and butorphanol (2.5 ml butorphanol 10 mg/ml) and were subsequently euthanized with an intracardiac injection of barbiturate. The liver was collected during necropsy and snap-frozen in liquid nitrogen, followed by storage at -80°C until liver transcriptomic analysis.

### RNA extraction and transcriptomic analysis

Liver tissues collected at euthanasia were used for transcriptomic analysis. RNA was extracted using the RNeasy Mini Kit (QIAGEN, USA), and whole-transcriptome shotgun sequencing was performed for gene expression profiling. Library preparation and sequencing were conducted by NOVOGENE (Cambridge, UK). RNA-Seq libraries were generated from 1000 ng of total RNA using the VAHTS mRNA-Seq V3 Library Prep Kit for Illumina (Vazyme, Nanjing, PRC) and sequenced on the Illumina NovaSeq 6000 platform, producing 150 bp paired-end reads. Raw reads underwent quality control and adapter trimming using TrimGalore (Babraham Bioinformatics, Cambridge, UK). The cleaned reads, averaging approximately 26 million per sample, were aligned to the porcine reference genome (Sscrofa11.1) using Tophat2. Gene annotation was obtained from Ensembl (release 99), and a gene-count matrix was generated using htseq-count.

For differential gene expression (DEG) analysis, weakly expressed genes (less than 10 copies in maximum 3 samples) were filtered out, and only protein-coding genes were included. DESeq2 package (version 1.38.3)^26,27^ was used to identify DEGs across three comparisons: survivors versus non-survivors, survivors versus uninfected controls, and non-survivors versus uninfected controls. Results were adjusted for sex and litter adjustment was performed for experiment 2. Fold-change values were estimated using the lfcShrink function with ashr, and an adjusted false discovery rate (FDR) of 0.05 was applied to determine statistically significantly enriched transcripts. Volcano plots depicting total gene expression were visualized using the EnhancedVolcano package (version 1.14.0)^28^.

Gene set enrichment analysis (GSEA) was performed to identify significantly altered biological pathways using the clusterProfiler package (version 4.0.2)^29^. Genes were ranked based on log2 fold-change values before pathway analysis. Pathway annotations were obtained from the Kyoto Encyclopedia of Genes and Genomes (KEGG) and the Biological Process (BP) aspect of Gene Ontology (GO) for *Sus scrofa*. An FDR threshold of 0.05 was used to define significantly enriched pathways. The normalized enrichment score (NES) was determined, with positive and negative values indicating upregulated and downregulated pathways, respectively. Heatmaps depicting the relative expression of genes within enriched pathways were generated on Z-score adjusted samples, using the ComplexHeatmap package (version 2.15.1)^30^.

### Cell culture experiments

The human monocyte-like THP-1 cell line was obtained from ATCC (TIB-202) and cultivated in RPMI-media (ThermoFisher 72400047) with 10% FCS (ThermoFisher A5670502). Prior to stimulation, cells were seeded in 24-well plates, 3 x 10^5^ cells/well and differentiated into macrophage-like cells by treatment with Phorbol-12-myristate-13-acetate (Sigma, 524400, 180 nM) for three days. Following differentiation, cells were infected with *S. epidermidis*, MOI 1 for 4 or 6 h, with and without treatment with BHB (1 mM), nicotinic acid (1 mM), oligomycin (1 uM; Sigma 75351) or MitoTempo (10 uM; Sigma, SML0737). Intracellular ATP was determined using the BacTiter Glow ATP assay (Promega, 74106) and cytokine secretion was determined using commercial ELISA kits (ThermoFisher 88-8086-86, 88-7106-86, 88-7066-86) according to manufacturer’s instructions. Total RNA was isolated using the RNeasy kit according to manufacturer’s instructions (Qiagen, 7404). cDNA was synthesized from µg of RNA using a High-Capacity cDNA Reverse Transcription Kit (Thermo Fisher, Waltham, MA, USA). Real-time quantitative PCR (RT–qPCR) was performed using the LightCycler 480 SYBR Green I Master kit on a LightCycler 480 (both Roche, Basel, Switzerland). Samples were analysed in duplicate using cytokine-specific primers. Gene expression was normalized to the expression of beta-actin using the 2^-△△CT^ method. Data was normalized to uninfected cells cultured in and all statistical comparisons between treatments were done by either paired T-test (for normally distributed data) or Wilcoxon’s signed rank test.

### Validation in child cohort

We validated the findings from piglets in children using data collected in the Copenhagen Prospective Studies on Asthma in Childhood 2010 (COPSAC_2010_) cohort, a Danish population-based birth cohort consisting of 700 mother-child pairs enrolled during pregnancy and followed prospectively through childhood. The cohort design, recruitment procedures, and baseline characteristics have been described in detail elsewhere^31^. Plasma samples were collected from healthy children at 18 months of age for metabolomic profiling, conducted by Metabolon, Inc. (Durham, NC, USA) using ultra-high-performance liquid chromatography-tandem mass spectrometry (LC-MS/MS). Details of the analytical procedures, including metabolite extraction, quality control, and compound identification, have likewise been described previously^32^. The metabolite data were semi-quantitative and included central ketone body metabolites. Data normalization and batch correction were performed by aligning median values and applying natural log transformation prior to statistical analysis. At the same timepoint, peripheral blood was collected in heparinized tubes and processed within four hours. The levels in pg/ml of the anti-inflammatory cytokine IL-10 and the pro-inflammatory cytokines interferon-gamma and IL-6 were determined in whole unstimulated blood using electrochemiluminescence-based multiplex assays (MesoScale Discovery) as described elsewhere ^29^.

### Statistics

Statistical analysis, data handling and visualization was performed in R (version 4.5) with the lme4, emmeans, ggplot2, dplyr, tidyr, and patchwork packages. For experiments containing only one litter of piglets, an additive linear model including Timepoint, Treatment Group and sex was fitted, and the main effects were evaluated by Type I ANOVA. At each timepoint, a separate linear model (Outcome ∼ Group + Sex + Birthweight) was fitted. Individual group differences within a timepoint were tested with estimated marginal means (emmeans) contrasts, using Tukey adjustment for multiple comparisons. If necessary, data were transformed (log, or square transformation) to ensure linearity. For experiments consisting of several litters, a linear mixed-effects model including the fixed factors Timepoint (3 h, 6 h, 12 h, Euthanasia), Treatment group (CON-SE, KETO-SE, RES-SE), sex and birthweight and the random intercept litter, was fitted. Fixed-effect significance was evaluated by F-tests with Satterthwaite degrees of freedom (lmerTest). If model assumptions were not met the outcome was transformed (log, inverse, or Box–Cox as appropriate) and the analysis repeated. At each timepoint a separate mixed model (Outcome ∼ Group + Sex + Birthweight + (1|Litter)) was fitted. Pair-wise group contrasts were obtained from estimated marginal means with Tukey adjustment for multiple testing. Clinical data from animal experiments are presented as truncated violin plots with line at median and depiction of individual replicates. In vitro experiments were analyzed using GraphPad prism 10.5.0. Typically, data were normalized to uninfected controls and compared using a ratio-paired t-test.

## Results

### Glycolytic inhibition attenuates inflammation and prevents lethal sepsis in piglets

We have previously shown that reducing parenteral glucose during neonatal infection decreases hepatic glycolysis and improves survival outcomes^15,17,18^. To assess the mechanistic role of glycolysis on sepsis pathogenesis, preterm piglets infected with live *S. epidermidis* were treated with the glycolysis inhibitor 2-DG and monitored for 20 h, or until humane endpoints (***Fig. 1A***). 2-DG treatment completely prevented lethal sepsis, with 100% survival in the treatment group compared to 55% in infected controls (***Fig. 1B***). Strikingly, blood bacterial density decreased over time in both groups, but were higher in the 2-DG-treated animals (***Fig. 1C***). Clinical parameters over the infection course were assessed via blood gas analysis. Blood pH was not significantly affected by 2-DG treatment (***Fig. 1D***). However, blood glucose levels were significantly higher in 2-DG-treated piglets compared to controls, with the difference decreasing over time driven by a decline in blood glucose levels in the 2-DG group (***Fig. 1E***). Lactate and partial pressure of CO were not significantly affected (**Fig. 1F, G**), while blood oxygen saturation was consistently higher in 2-DG-treated animals throughout the experiment (***Fig. 1H***). Hematological analysis revealed a typical decline in leukocyte and neutrophil counts in all infected animals over time. Interestingly, 2-DG treatment led to lower leukocyte and lymphocyte counts at 6- and 12-h post-infection and lower erythrocyte counts over the course of the experiment, relative to the infected controls (***Fig. S1A-F***).

**Figure 1:**
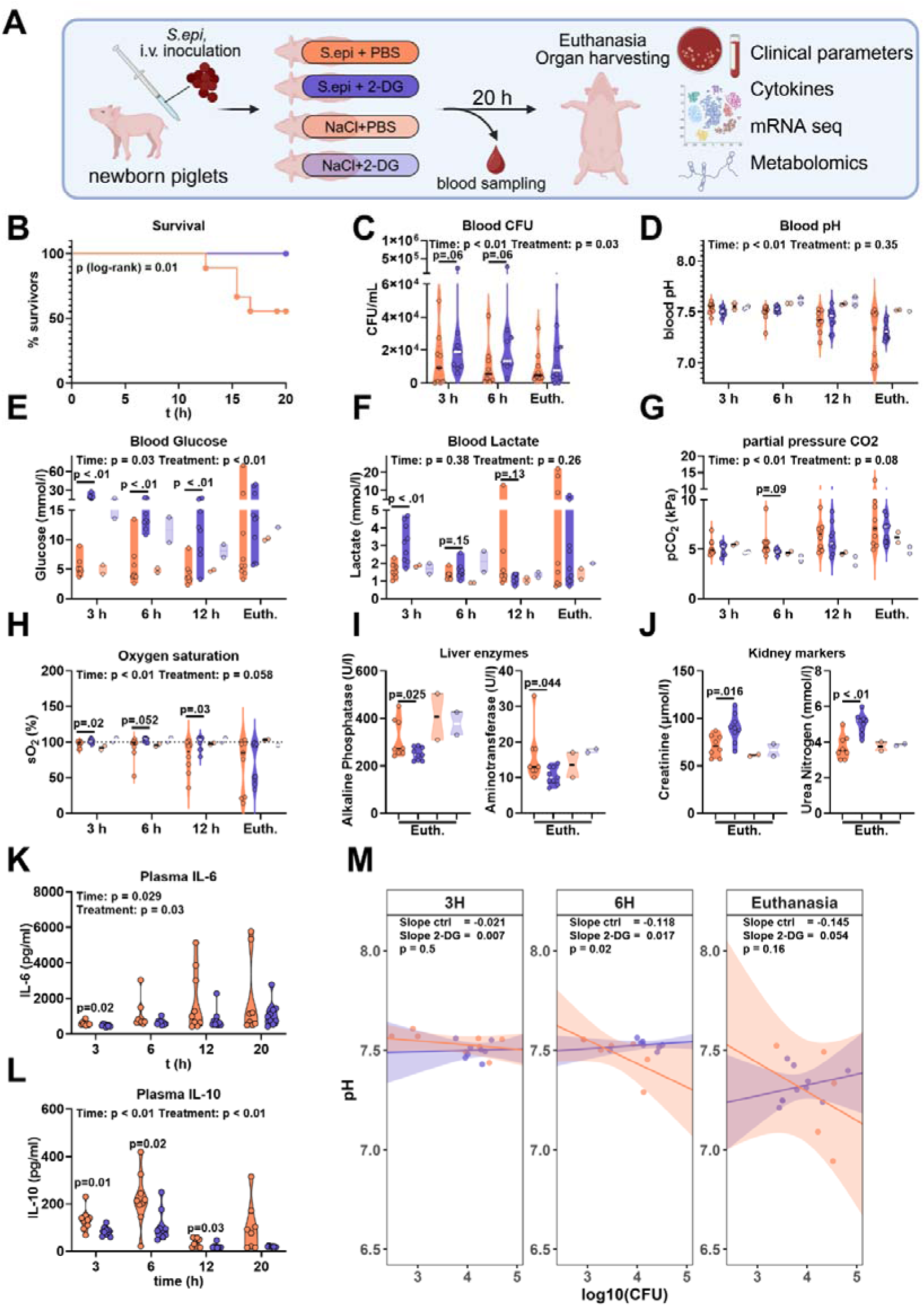
Clinical effects of 2-DG supplementation on sepsis pathogenesis in infected piglets. **A)** Experimental setup: preterm neonatal piglets were infected with *S. epidermidis*, 10^9^ CFU/kg with or without treatment with 2-DG, 750 mg/kg. Animals were cared for for 20 h or until humane euthanasia. **B)** Survival, presented as time to euthanasia according to defined humane endpoints. Depicted are infected controls and infected 2-DG treated animals. **B)** Blood bacterial counts of infected animals, depicted as CFU/ml. **D-H)** Blood gas parameters. **D)** Blood pH. **E)** Blood glucose in mmol/l. **F)** Blood lactate in mmol/l. **G)** Partial pressure of CO_2_ (kPA). **H)** Blood oxygen saturation in %. **I, J)** Biochemical analysis of blood plasma at euthanasia. **I)** Activity of alkaline phosphatase and alanine-aminotransferase in plasma, depicted in U/l. **J)** Blood plasma levels of Creatinine (µmol/l) and urea nitrogene (mmol/l). **K, L)** Blood plasma levels of IL-6 **(K)** and IL-10 **(L)** in pg/ml, measured by ELISA. **M)** Resistance-tolerance-analysis. A reaction-norm-analysis was performed to plot blood pH was plotted against the log_10_(CFU) and slopes calculated by linear regression. Slopes were compared using a sum-of-squares F-test. N = 10-12 (infected groups); 2 (control groups); line at median.

At euthanasia, plasma levels of the liver enzymes alkaline phosphatase and aminotransferase were significantly lower in 2-DG-treated pigs, indicating 2-DG treatment protected against sepsis-induced liver injury (***Fig. 1I***). In contrast, creatinine and urea nitrogen levels were significantly increased in response to 2-DG, which may relate to an increased protein catabolism during inhibition of glycolysis. Further, 2-DG also significantly reduced levels of plasma cytokines IL-6 and IL-10 (***Fig. 1K-L***) but did not affect TNF-α levels (***Fig. S1G***). The TNF-α/IL-10 ratio, a marker of pro-versus anti-inflammatory balance, remained unchanged (***Fig. S1H***). We plotted blood pH against bacterial burden to assess disease tolerance, defined as the ability to maintain health with increasing pathogen burden (***Fig. 1M***). A steeper negative slope indicates reduced tolerance, as pH drops more sharply with increasing bacterial load. Over time, infected controls developed steeper slopes, reflecting worsening physiological status with rising pathogen levels. In contrast, 2-DG–treated animals maintained shallower slopes, indicating improved tolerance. Thus, 2-DG appeared to enhance disease tolerance by preserving systemic stability despite infection.

Taken together, these results demonstrate that glycolytic inhibition prevents lethal sepsis in infected piglets and indicate that this occurs through modulating the inflammatory response and systemic metabolism and shifting the defense strategy from resistance toward tolerance.

### Glycolytic inhibition reduces hepatic inflammation and rewires hepatic metabolism in infected piglets

Next, we performed RNA sequencing of liver tissue to investigate transcriptomic changes induced by 2-DG treatment.

GSEA-GO revealed a significant inhibition of immune-related pathways in 2-DG-treated piglets compared to infected controls (***Fig. 2A***). Further KEGG pathway analysis showed a metabolic shift, with upregulation of alternative energy metabolism pathways, including oxidative phosphorylation, indicating a reprogramming of energy metabolism away from glycolysis (***Fig. 2B***).

**Figure 2:**
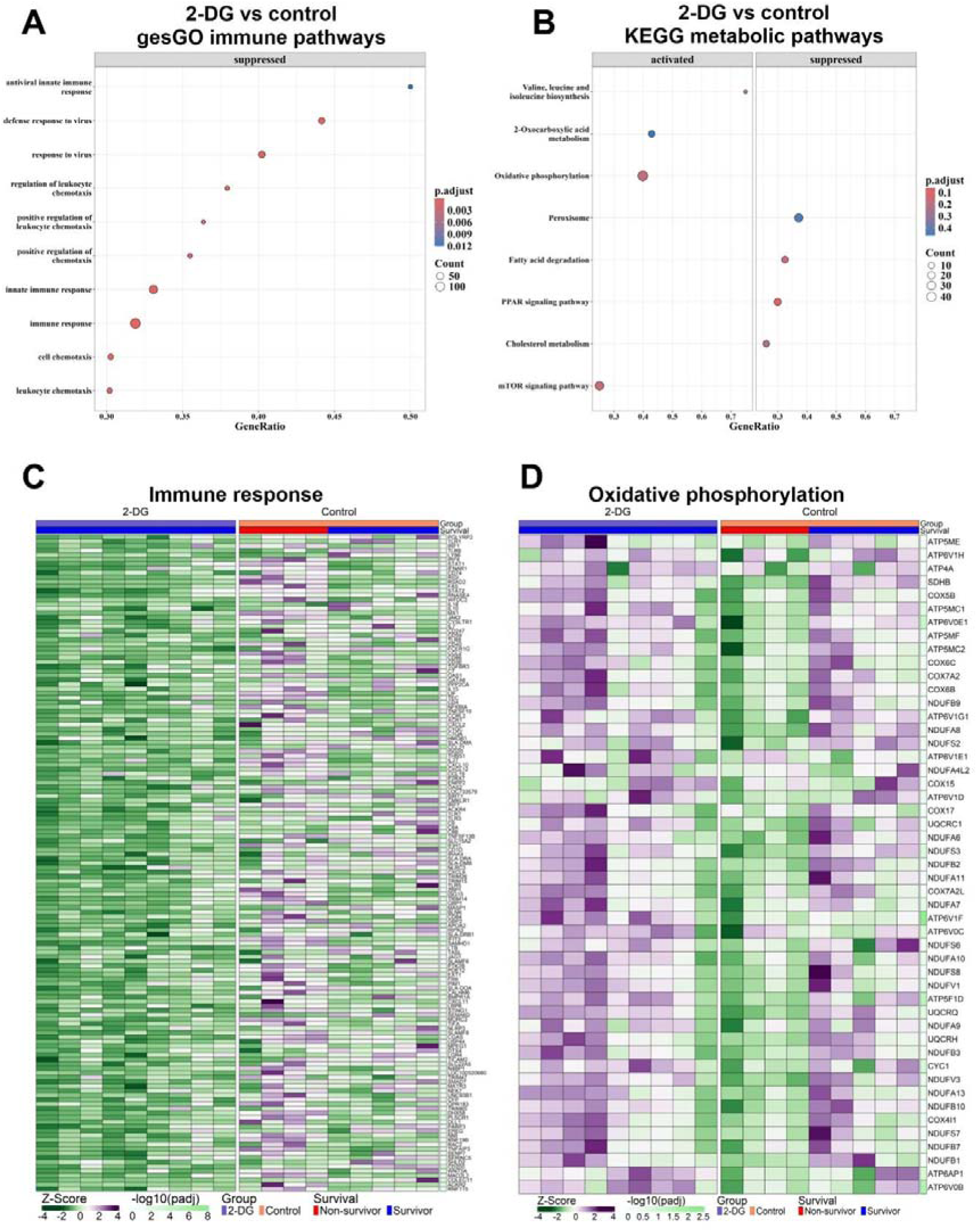
Impact of 2-DG treatment on the hepatic transcriptome during infection in piglets. **A)** Gene-set enrichment analysis (GSEA) according to the gene ontology (GO) database (biological pathway), comparing 2-DG treated, infected piglets vs infected controls. Immune-associated pathways were selected manually. The labels “activated” and “suppressed” indicate direction of regulation in 2-DG group. **B)** GSEA according to the KEGG database. Metabolism-associated pathways were selected manually. **C, D)** Heatmap of genes associated with the pathways “Immune Response (GO-BP)” **(C)** and “Oxidative phosphorylation (KEGG)” **(D)**. Expression is depicted as Z-scores, calculated based on all samples. Depicted p-values are calculated between 2-DG, infected piglets and infected controls. N = 10-12 (infected groups); 2 (control groups).

Immune response-associated genes were strongly downregulated in 2-DG-treated piglets compared to controls. Interestingly, within the control group, survivors exhibited a lower expression of immune-related genes compared to non-survivors, suggesting a link between immune activation and disease severity (***Fig. 2C***). A similar effect was observed when investigating TNF-α signaling associated genes (***Fig. S2A***). By contrast, OXPHOS-associated genes were upregulated in 2-DG treated piglets and, within the control group, OXPHOS was reduced in non-survivors, compared to survivors ***(Fig. 2D)***. We further analyzed mitochondrial function by examining the expression of genes encoding mitochondrial protein complexes. GO-CC analysis revealed an activation of genes associated with the mitochondrial compartment upon 2-DG treatment (***Fig. S2C***). Mitochondrial activity increased upon 2-DG treatment, supporting the hypothesis of altered energy metabolism as a mediator of infection survival (***Fig. S2B, D***). Interestingly, survivors also showed markedly enhanced mitochondrial activity compared to non-survivors (***Fig. S2B, D***). Overall, 2-DG-treated piglets had gene expression patterns like surviving controls, suggesting a protective metabolic shift. In summary, 2-DG treatment reduced inflammation and activated mitochondria-associated metabolic pathways, in particular oxidative phosphorylation. These findings suggest that 2-DG treatment reprograms metabolism and suppresses immune activation, leading to the observed disease tolerance phenotype; contributing to improved survival at the cost of pathogen proliferation in the infected piglets.

### Beta-hydroxybutyrate inhibits inflammation and abolishes sepsis mortality

Based on the 2-DG results obtained, we reasoned that boosting OXPHOS and mitochondrial metabolism during neonatal infection could be a promising strategy to improve disease tolerance phenotypes and improve infection outcomes. Since BHB can serve as a fuel source for OXPHOS, and has been shown to improve mitochondrial function, we hypothesized that a partial substitution of glucose with BHB in the piglet sepsis model improves sepsis outcomes. To test this, we aimed to develop a BHB supplementation protocol that satisfies neonatal energy demands without inducing hyperglycemia and sepsis. Infected preterm piglets received PN containing either 10% glucose (as in the 2-DG experiment), 2.5% glucose, or 2.5% glucose with 2.5% BHB (***Fig. 3A***). Notably, BHB supplementation completely prevented lethal sepsis, with 100% survival in the ketone-treated group compared to 45% in the 10% glucose control group and 80% in the restricted glucose group (***Fig. 3B***). Bacterial counts did not differ among the three infected groups (***Fig. 3C***). Blood gas analysis showed that BHB treatment completely prevented blood acidosis whereas the remaining two groups started to become acidotic 12 h post-bacterial challenge (***Fig. 3D***). Likewise, standard bicarbonate levels (***Fig. 3H***) and base excess (***Fig. S3A***) were significantly increased by BHB treatment across all time points, all indicating improved acid-base balance. Blood glucose levels were significantly reduced when comparing the BHB group to the 10% glucose group but not affected when compared to 2.5% glucose treatment. Notably, glucose reduction alone did not significantly affect acid-base balance. While oxygen saturation decreased over time in all groups, it was significantly improved by BHB supplementation after 12 h (***Fig. 3G***). Hematological analysis showed a typical decline in leukocyte and platelet counts in all infected animals over time, with no significant effect of BHB treatment (***Fig. S3B-F***).

**Figure 3:**
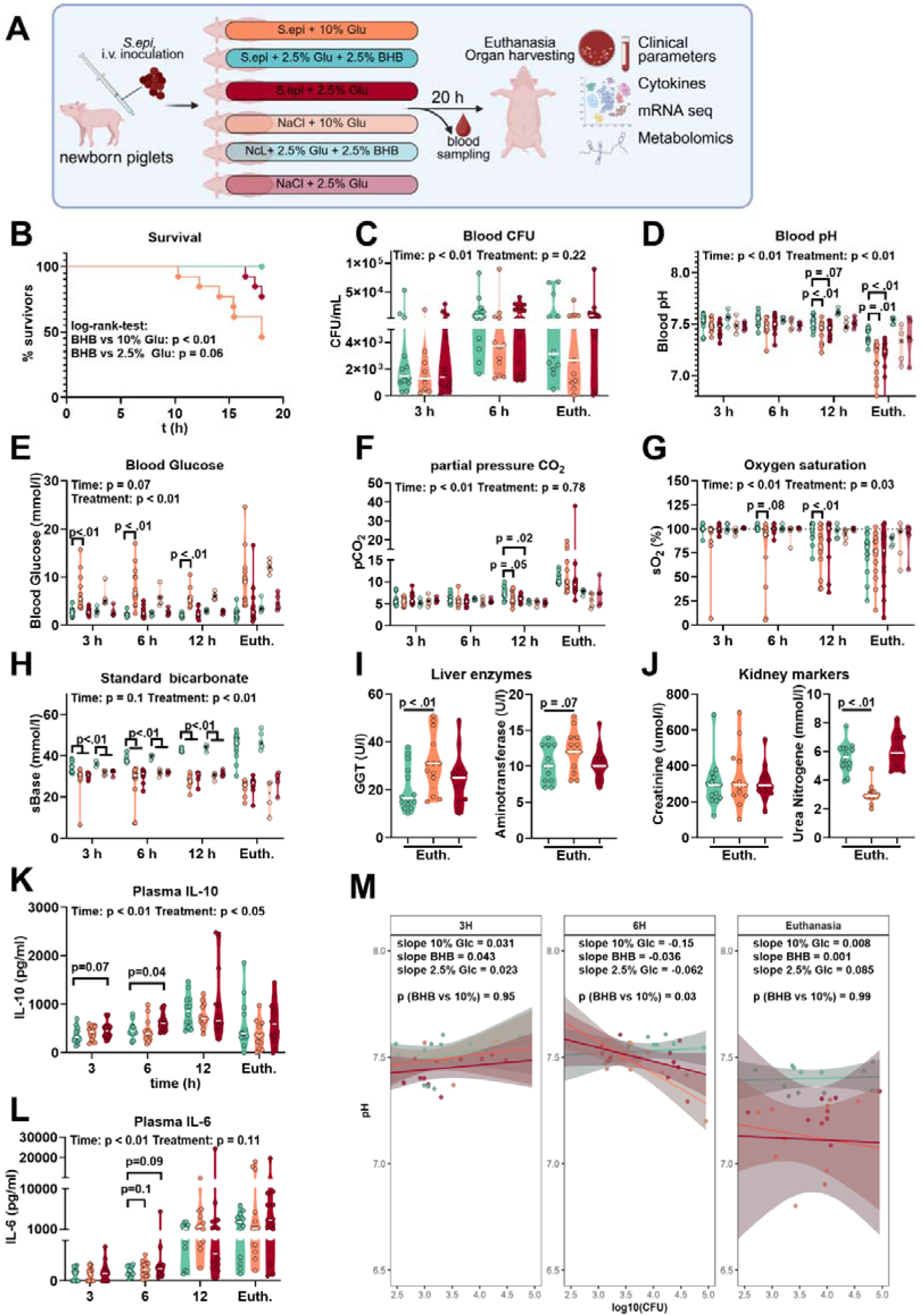
Clinical effects of BHB supplementation on sepsis pathogenesis in infected piglets. **A)** Experimental setup: preterm neonatal piglets were infected with *S. epidermidis*, 10^9^ CFU/kg. Controls were left uninfected. Over the course of the experiment, piglets received PN containing either 10% Glucose, 2.5% Glucose or 2.5% Glucose and 2.5% BHB. Animals were cared for for 20 h or until humane euthanasia. **B)** Survival, presented as time to euthanasia according to defined humane endpoints. Only infected groups are presented. **C)** Blood bacterial counts of infected animals, depicted as CFU/ml. Blood samples taken at indicated timepoints were diluted in PBS and plated on blood agar plates. **D-H)** Blood gas parameters. **D)** Blood pH. **E)** Blood glucose in mmol/l. **F)** Partial pressure of CO_2_ (kPA). **G)** Blood oxygen saturation in %. **H)** Standard bicarbonate in mmol/l. **I, J)** Biochemical analysis of blood plasma at euthanasia. **I)** Activity of gamma-glutanyl-transferase and alanine-aminotransferase in plasma, depicted in U/l. **J)** Blood plasma levels of Creatinine (µmol/l) and urea nitrogene (mmol/l). **K, L)** Blood plasma levels of IL-6 **(K)** and IL-10 **(L)** in pg/ml, measured by ELISA. **M)** Resistance-tolerance-analysis. A reaction-norm-analysis was performed to plot blood pH against the log_10_(CFU), with slopes calculated by linear regression. Slopes were compared using a sum-of-squares F-test. N = 13-15 (infected groups); 4-5 (control groups); line depicts the median.

Organ function was assessed via biochemical analysis of blood plasma at euthanasia. Plasma levels of the liver enzymes alanine aminotransferase and gamma-glutamyl transferase were significantly reduced in the BHB-treated group, indicating less liver injury (***Fig. 3I***). Again, glucose reduction alone did not show significant effects. Additionally, increased plasma urea nitrogen levels, without corresponding increase in creatinine, suggested increases in amino acid catabolism in response to BHB supplementation (***Fig. 3J***). Cytokine analysis revealed that BHB treatment significantly or by tendency reduced plasma cytokine levels at early phase of infection (***Fig. 3K-L***), supporting a potential anti-inflammatory effect. Reaction norm analysis, indicating the state of disease tolerance, further demonstrated that BHB-treated piglets exhibited a shallower slope after 6 h, in contrast to the steeper slopes observed in the 10% and 2.5% glucose groups. At euthanasia, blood pH values in the control groups stabilized at a low level, indicative of declined health, whereas this effect was absent in ketone-treated piglets (***Fig. 3M***).

In summary, BHB treatment during bloodstream infection in preterm pigs significantly improved survival by supporting acid-base balance and boosting anti-inflammatory and disease tolerance responses.

### BHB treatment attenuates hepatic inflammation

To investigate whether alterations in the host energy metabolism may be responsible for the sepsis-protective effect of BHB, we performed an RNAseq analysis of the hepatic transcriptome. A GSEA-GO analysis revealed a suppression of several inflammation-associated pathways in BHB treated animals (innate immune response, defense response; p<0.01), compared to the restricted glucose group (***Fig. 4A*, *C***). This confirms the general anti-inflammatory function of BHB. Compared to the group receiving 10% glucose, we further found BHB treatment to activate pathways associated with mitochondrial metabolism (organic acid metabolic process), growth and development (embryonic morphogenesis) (***Fig. 4B, D***), indicating a metabolic rewiring. These pathways involve the biosynthesis of short-chain fatty acids such as propionate and methylmalonate and may be involved in anti-inflammatory responses. Related pathways were previously reported to be associated with sepsis survival ^16^. Moreover, we have previously shown that these pathways are important for both infection survival and maintenance of disease tolerance in the same animal model. In summary, we found the sepsis protective effects of BHB to be accompanied by reduced inflammation and metabolic rewiring in hepatic tissues.

**Figure 4:**
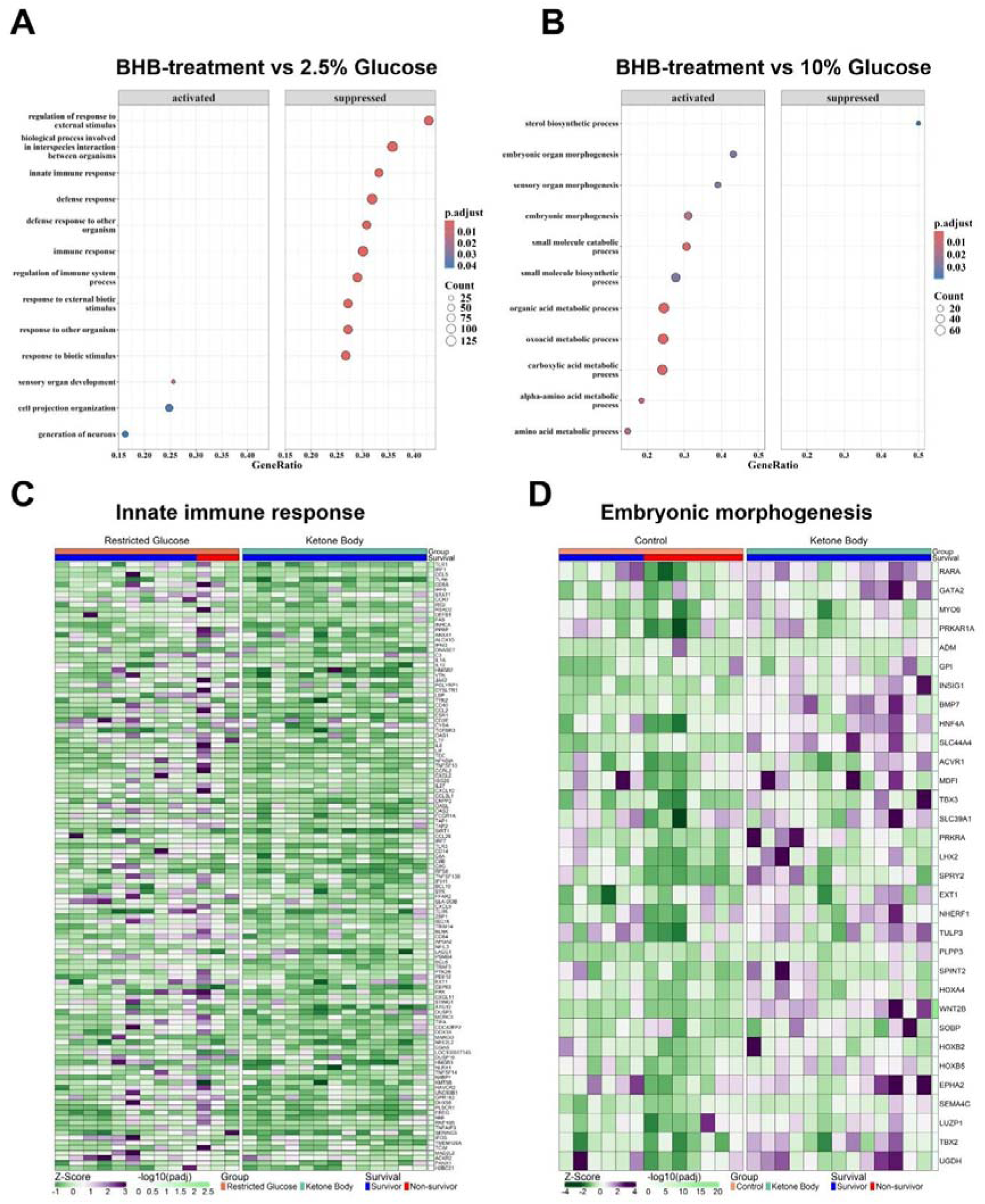
Impact of BHB supplementation on the hepatic transcriptome during infection in piglets. **A)** GSEA-GO database (biological pathway) analysis, comparing BHB+2.5% Glucose-treated, infected piglets vs 2.5% Glucose/infected. Immune-associated pathways were selected manually. **B)** GSEA according to the KEGG database. Metabolism-associated pathways were selected manually. The labels “activated” and “suppressed” indicate direction of regulation in BHB group. **C, D)** Heatmap of genes associated with the pathways “Immune Response (GO-BP)” **(C)** and “Oxidative phosphorylation (KEGG)” **(D)**. Expression is depicted as Z-scores, calculated based on all samples. Depicted p-values are calculated between 2-DG, infected piglets and infected controls. N = 13-15 (infected groups); 4-5 (control groups).

### BHB treatment during infection inhibits inflammation *in vitro*

Next, we aimed to investigate whether BHB can induce an anti-inflammatory response in human cell cultures. THP-1 macrophages were infected with *S. epidermidis*, MOI1 for 6 h, with and without treatment with 1 mM BHB. Gene expression analysis by qPCR revealed that BHB treatment during infection significantly reduced the expression of TNF-α, IL-6, and IL-8, cytokines associated with the NF-κB signaling pathway (***Fig. 5A***). This aligns with hepatic RNAseq results in infected piglets, where NF-κB associated pathways were inhibited by BHB treatment. We further found BHB treatment to increase intracellular levels of ATP, indicating an elevated OXPHOS (***Fig. 5B***). Interestingly, BHB treatment during stimulation with heat-inactivated bacteria or treatment with the bacterial ligands lipoteichoic acid (LTA) or lipopolysaccharide (LPS) had no or minor effects on inflammation (***Fig. 5C, D***).

**Figure 5:**
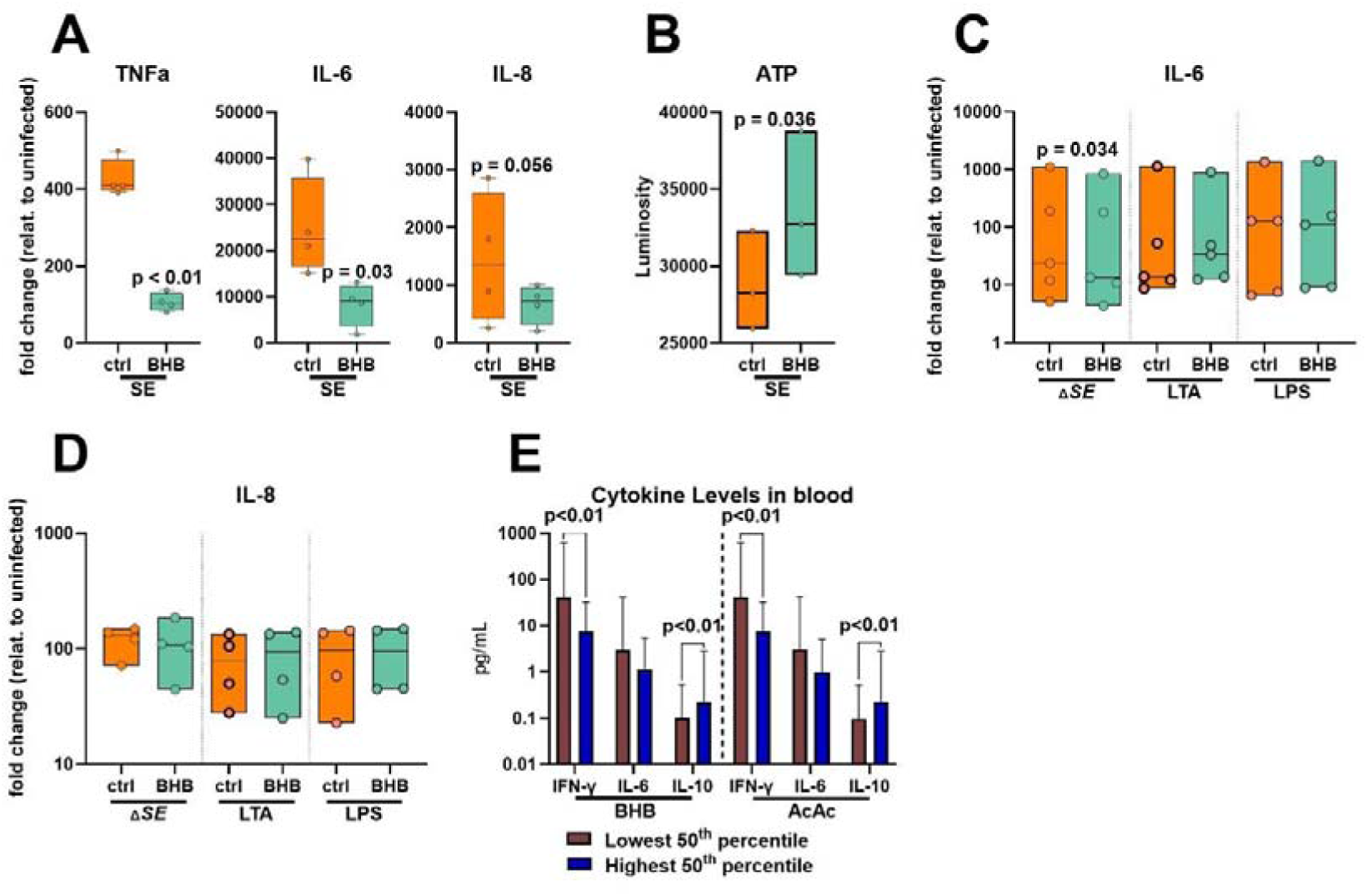
BHB inhibits inflammatory cytokines and increases intracellular ATP in infected human macrophages and children. **A, B)** PMA-differentiated THP-1 cells were infected with *S. epidermidis*, MOI1, for 6 h with or without treatment with 1 mM BHB. **A)** Expression of indicated cytokines, depicted as fold changes compared to uninfected controls. **B)** ATP, measured by bioluminescence, depicted as fold changes compared to uninfected controls. **C, D)** Differentiated THP-1 cells were treated with heat-inactivated *S. epidermidis* (MOI 100), LTA (10 µg/ml), LPS (100 ng/ml) with and without for 6h, with and without BHB treatment. Expression of indicated cytokines was determined by qPCR. **E)** Blood plasma levels of BHB and Acetoacetate (AcAc) and indicated cytokines were measured in blood of 538 18-month-old children. Cytokine levels were separated into two groups based on lowest and highest 50% levels of BHB and AcAc, respectively. N=3-5 (A-D).

### Plasma BHB levels are associated with reduced inflammatory responses in children

Finally, we aimed to confirm the relationship between BHB and anti-inflammatory responses in children. Blood and plasma samples were collected from 538 of COPSAC_2010_-enrolled children at 18 months of age and subjected to metabolomics profiling and cytokine measurement. Children with the highest average levels of the two ketone bodies (BHB and Acetoacetate) exhibited significantly higher levels of the anti-inflammatory cytokine IL-10, and lower levels of the pro-inflammatory cytokine interferon-gamma. Moreover, they trended to display lower levels of IL-6 (Fig 5E). Thus, these human data from a birth cohort confirms an anti-inflammatory role of BHB observed in the piglet experiments.

## Discussion

In this study, we demonstrated that metabolic modulation through glycolytic inhibition or ketone body supplementation significantly improves neonatal sepsis outcomes in infected preterm piglets. Strikingly, these protective effects are associated with enhanced disease tolerance, reduced systemic inflammation and liver injury. These results underscore metabolic rewiring as a promising non-antimicrobial therapeutic strategy for neonatal infection.

Effective host defense strategies against infection rely on balancing disease resistance, characterized by pathogen elimination, and disease tolerance, which limits tissue damage without necessarily reducing pathogen load. Resistance is energetically demanding and typically associated with glycolytic metabolism and pro-inflammatory responses, whereas tolerance mechanisms rely more heavily on mitochondrial OXPHOS and anti-inflammatory pathways^3,15,33^. Given their limited energy reserves and the need to prioritize energy for growth and development, infants are less capable of sustaining energetically costly inflammatory responses ^3^. As a result, they are more likely to adopt a disease tolerance strategy during early phases of infection. However, circulating bacteria may reach critical levels, forcing the host’ immune system to switch to disease resistance, now with excessive glycolysis-induced inflammatory responses to high amounts of pathogens, causing collateral tissue damage. We reason that metabolic interventions that maintain mitochondrial metabolism and prevent glycolytic switch may balance the two defense strategies and improve infection outcomes. In this study, we provide strong evidence supporting these with 2-DG and BHB interventions in infected piglets.

Glycolytic inhibition with 2-DG resulted in reduced inflammation, improved acid-base balance, enhanced mitochondrial activity, and complete prevention of lethal sepsis. This strongly supports previous findings that excessive glycolysis drives inflammation and organ dysfunction in neonatal sepsis^15,17,34^. Previously, 2-DG has been shown to likewise inhibit immune responses during sterile inflammation in mice^35^. During disease resistance, the host responds to bacterial invasion by mounting a pro-inflammatory reaction. When this becomes excessive, it can be accompanied by a drop in blood pH, which becomes more pronounced as bacterial burden increases. When plotted, this relationship manifests as a negative slope between blood pH and bacterial burden^15,36^. In contrast, disease tolerance is indicated by a shallow slope, as increases in bacterial burdens do not affect the clinical response. Using this analysis, we found 2-DG treatment to induce disease tolerance in our piglet model of sepsis. Further analysis of hepatic tissues clarified how this metabolic rewiring mediated the shift. Inhibition of hepatic glycolysis activated alternative energy pathways, including OXPHOS and citrate cycle activity, while concurrently suppressing inflammatory pathways. Notably, the hepatic transcriptional profile induced by 2-DG closely resembled that of surviving controls, supporting metabolic reprogramming as key to enhanced survival outcomes. This aligns with prior studies emphasizing the liver as a central metabolic and immune regulator in sepsis as well as being key for the establishment of disease tolerance^8,37,38^. Strikingly, 2-DG treated animals had higher bacterial burdens than the infected controls, but they could still gradually clear bacteria over the course of infection. This suggests that blocking glycolysis and enhancing OXPHOS do not block disease resistance response but rather maintain a sufficient balance between the two disease strategies to survive infection.

Building on these insights, we explored ketone body supplementation as a nutritional strategy to alter hepatic energy metabolism. Ketones such as BHB serve as efficient mitochondrial fuels, boosting OXPHOS and reducing reactive oxygen species production^39^. Our data showed that partial replacement of glucose with BHB in PN for infected preterm pigs fully prevented sepsis mortality and improved clinical parameters such as blood pH, acid-base balance, oxygen saturation, and liver function. Importantly, while a ketogenic diet was sometimes associated with increased risks for kidney damage^40^, we found no effect of BHB treatment on kidney markers in plasma. BHB treatment was likewise associated with improved disease tolerance, indicated by a stable blood pH over the course of infection. However, despite improved tolerance, the pathogen burden did not differ between treatment groups, underscoring that the intervention enhanced disease tolerance without compromising antibacterial defenses. Previous studies have reported similar anti-inflammatory and organ-protective effects of ketogenic diets in diseases like epilepsy, autoimmune conditions, and viral infections ^41–44^. Mechanistically, BHB treatment also inhibited hepatic inflammatory pathways associated with NF-kB signaling. Cell culture experiments with THP-1 derived macrophages confirmed these anti-inflammatory effects. Thus, BHB’s protective mechanisms may involve dual roles as both metabolic fuel and signaling molecules capable of directly attenuating inflammatory responses to infections. We lastly validated our findings in a human context through analysis of metabolomics and cytokine profiles from healthy 18-month-old infants. Here, the group of children with the highest plasma BHB levels displayed the lowest levels of pro-inflammatory interferon-gamma and highest levels of the anti-inflammatory cytokine IL-10, consistent with the anti-inflammatory effects of BHB demonstrated in our animal and in vitro cellular data. It must be noted that this correlation was observed in unstimulated *ex vivo* samples, of young children, not infants. Still this alignment between the piglet and human data strengthens the translational validity of our animal-model observations, indicating that BHB may modulate pro-inflammatory responses similarly in humans, also beyond the neonatal period.

Neonatal sepsis remains challenging to manage due to the limitations of antimicrobial strategies and the underdeveloped neonatal immune system. Our results support metabolic rewiring using ketone supplementation, either alone or as part of PN solutions, as an innovative approach to improve defense strategies and neonatal infection outcomes, particularly for preterm infants routinely exposed to glucose-rich PN. The translational potential of our findings is substantial, particularly given the widespread clinical use and safety profile of ketone supplements^45–48^.

There are limitations to our study. First, our findings are based on a neonatal piglet model. While this model closely mirrors neonatal immune responses and sepsis pathophysiology^49^, species-specific responses cannot be entirely ruled out. Additionally, using BHB chloride resulted in artificially elevated sodium levels, which may not be desirable clinically. Thus, this study can only provide proof of concept, and further investigation into alternative ketone formulations, such as ketone esters or HCAR2-specific agonists, is warranted.

## Supplemental figures

**Supplemental Figure 1:**
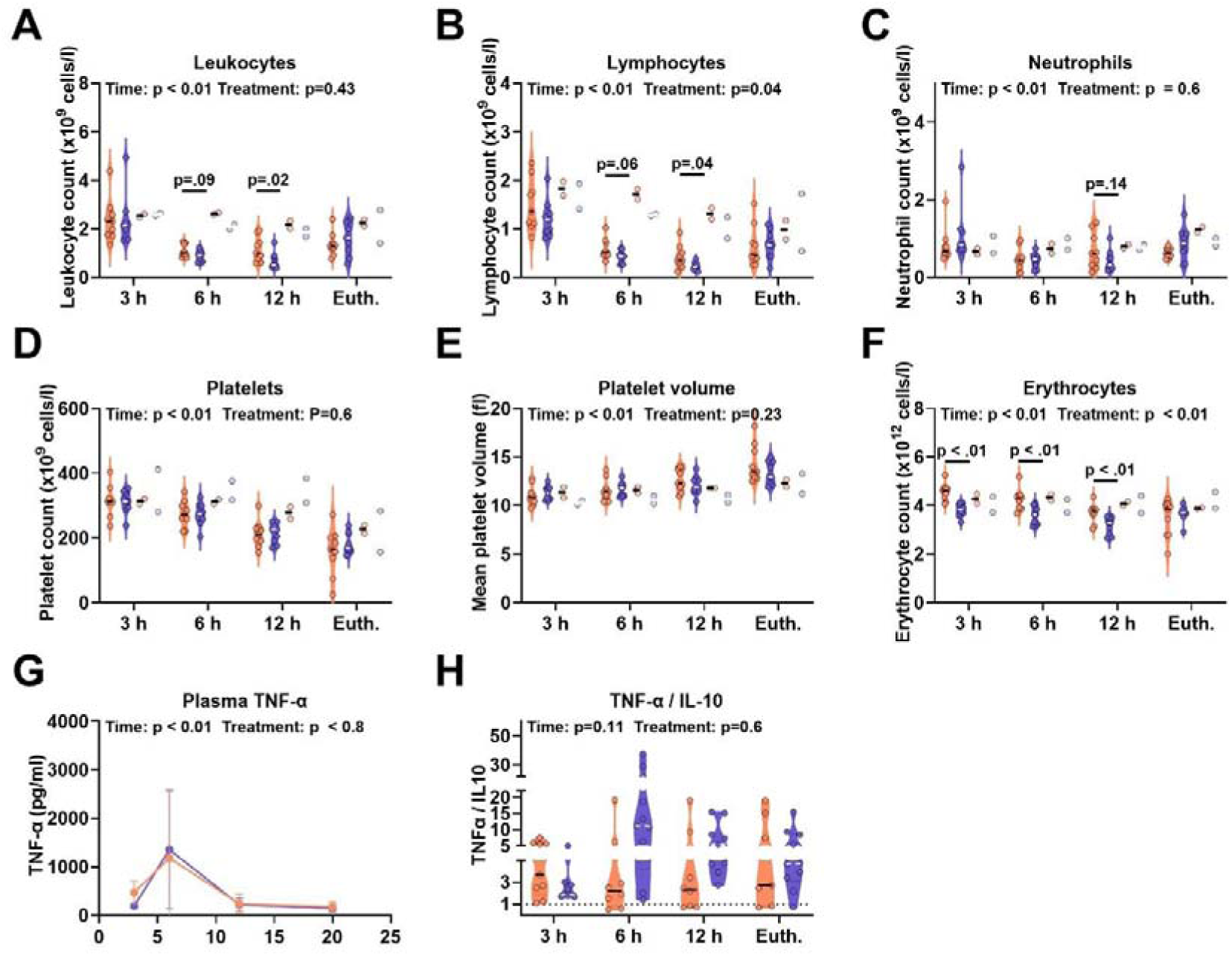
Extended clinical data following 2-DG treatment. **A-F)** Hematological analysis of blood samples taken at indicated timepoints following infection. **A)** Total Leukocytes. **B)** Total lymphocytes. **C)** Neutrophil counts. **D)** Platelet counts. **E)** Mean platelet volume. **F)** Erythrocytes. G) TNF-α levels in plasma, measured by ELISA at indicated timepoints. **H)** Quotient TNF-α/IL-10, measured by ELISA, at indicated timepoints. **Statistics:** Statistical analysis was performed in R (version 4.x) with the lme4, emmeans, ggplot2, dplyr, tidyr, and patchwork packages. An additive linear model including Timepoint (3 h, 6 h, 12 h, Euthanasia), Treatment Group (CON-SE vs 2DG-SE) and sex was fitted, and the main effects were evaluated by Type I ANOVA. At each timepoint, a separate linear model (Outcome ∼ Group + Sex + Birthweight) was fitted. Individual group differences within a timepoint were tested with estimated marginal means (emmeans) contrasts, using Tukey adjustment for multiple comparisons. If necessary, data were transformed (log-or square transformation) to ensure linearity. N = 10-12 (infected groups); 2 (control groups); line at median.

**Supplemental Figure 2:**
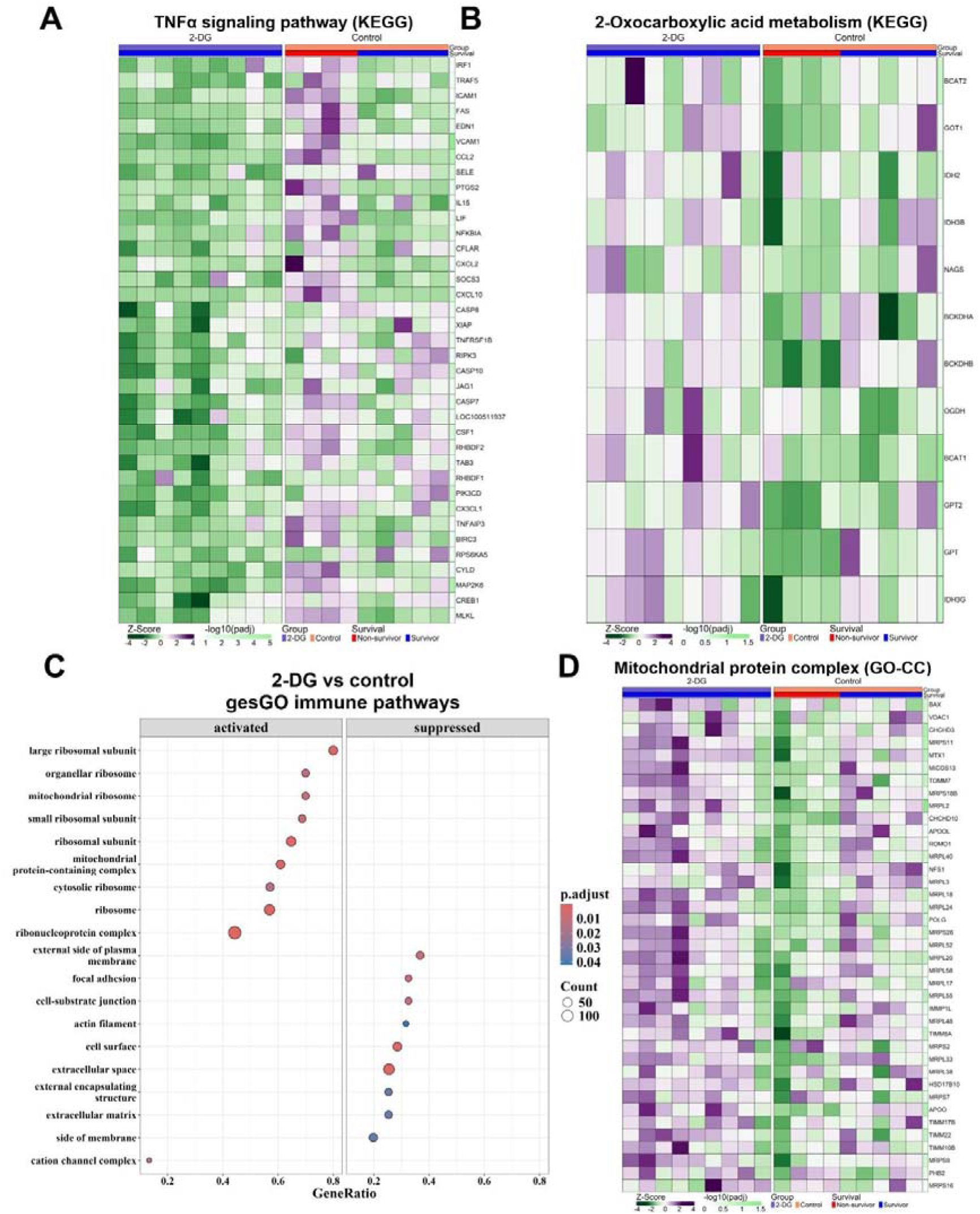
Impact of 2-DG treatment on the hepatic transcriptome during infection. **A, B)** Heatmaps of genes associated with the KEGG-pathways “TNFα signaling pathway” **(A)** and “2-Oxocarboxylic acid metabolism” **(B). C)** GSEA according to the GO database (cellular compartment), comparing 2-DG treated, infected piglets vs infected controls. **D)** Heatmap associated with the GO-CC pathway “Mitochondrial protein complex”. Gene expression is depicted as Z-scores, calculated based on all samples. Depicted p-Values are calculated between 2-DG, infected piglets and infected controls. N = 10-12 (infected groups); 2 (control groups); line at median.

**Supplemental Figure 3:**
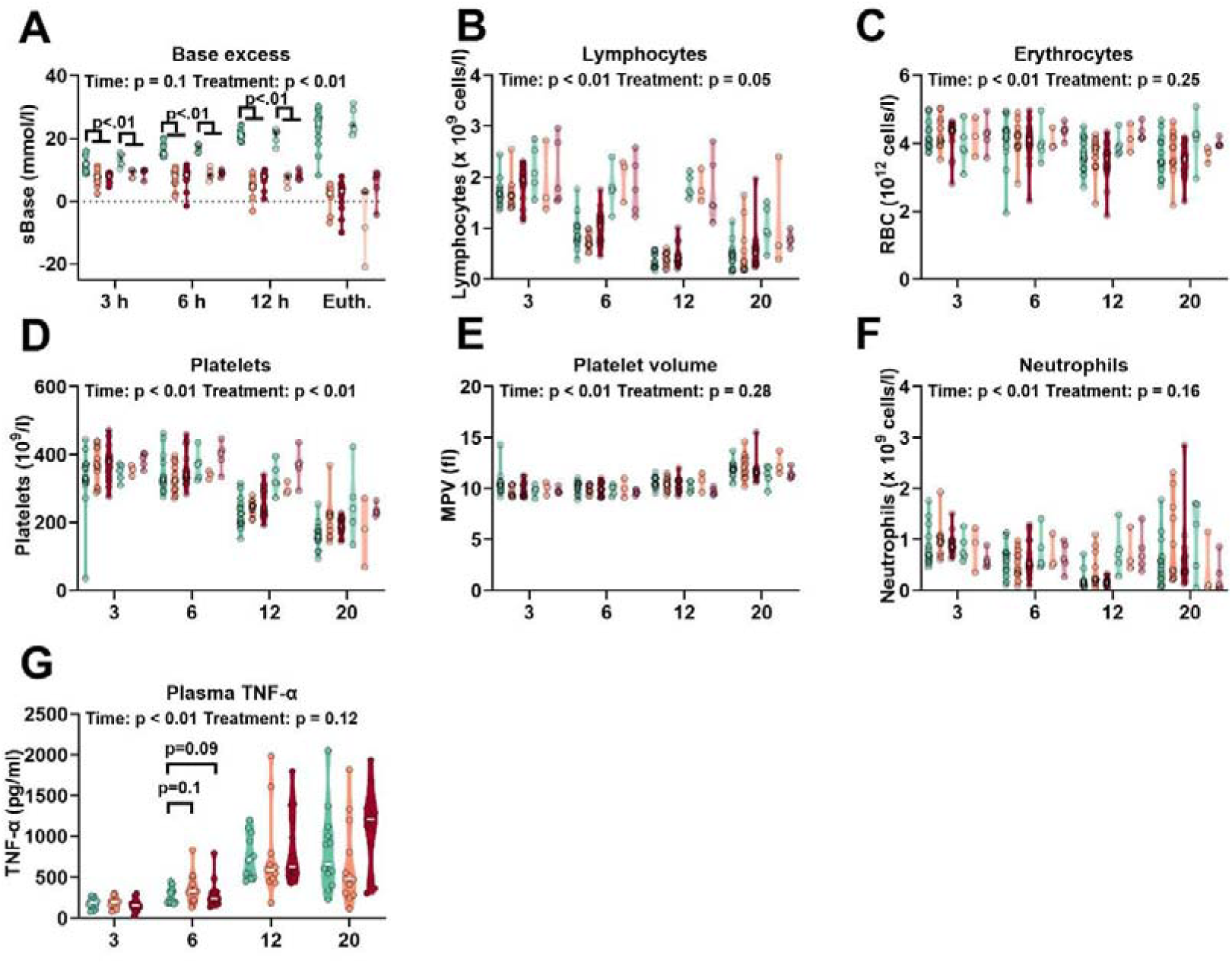
Extended clinical data following BHB treatment. **A)** Base excess in whole blood, measured by blood gas analysis at indicated timepoints. **B-F)** Hematological analysis of samples taken at indicated timepoints following infection. **B)** Total lymphocytes. **C)** Total erythrocytes. **D)** Total platelets. **E)** Mean platelet volume. **F)** Total neutrophils. **G)** Blood plasma levels of TNFα at indicated timepoints, measured by ELISA. Statistical analysis was performed in R (v 4.x) with the lmerTest, lme4 and emmeans packages. For each variable a linear mixed-effects model including the fixed factors Timepoint (3 h, 6 h, 12 h, Euthanasia), Treatment group (CON-SE, KETO-SE, RES-SE), birthweight and sex and the random intercept Litter, was fitted. Fixed-effect significance was evaluated by F-tests with Satterthwaite degrees of freedom (lmerTest). If model assumptions were not met the outcome was transformed (log, inverse, or Box–Cox as appropriate) and the analysis repeated. At each timepoint a separate mixed model (Outcome ∼ Group + Sex + (1|Litter)) was fitted. Pair-wise group contrasts were obtained from estimated marginal means with Tukey adjustment for multiple testing. N = 13-15 (infected groups); 4-5 (control groups); line at median.

